# Latent circuit inference from heterogeneous neural responses during cognitive tasks

**DOI:** 10.1101/2022.01.23.477431

**Authors:** Christopher Langdon, Tatiana A. Engel

## Abstract

Higher cortical areas carry a wide range of sensory, cognitive, and motor signals supporting complex goal-directed behavior. These signals are mixed in heterogeneous responses of single neurons tuned to multiple task variables. Dimensionality reduction methods used to analyze neural responses rely merely on correlations, leaving unknown how heterogeneous neural activity arises from connectivity to drive behavior. Here we present a framework for inferring a low-dimensional connectivity structure—the latent circuit—from high-dimensional neural response data. The latent circuit captures mechanistic interactions between task variables and their mixed representations in single neurons. We apply the latent circuit inference to recurrent neural networks trained to perform a context-dependent decision-making task and find a suppression mechanism in which contextual representations inhibit irrelevant sensory responses. We validate this mechanism by confirming the behavioral effects of patterned connectivity perturbations predicted by the latent circuit structure. Our approach can reveal interpretable and causally testable circuit mechanisms from heterogeneous neural responses during cognitive tasks.

---

Cognitive functions depend on higher cortical areas, which integrate diverse sensory and contextual signals to produce a coherent behavioral response. These neural signals arise from excitatory and inhibitory interactions in cortical circuits. Traditionally, hand-crafted neural circuit models were used to pose specific mechanistic hypotheses about how behavioral responses arise from excitation and inhibition between a few neural populations representing task variables^1–11^. By linking connectivity, activity, and function, these models can predict changes in behavioral performance under perturbations of the circuit structure (e.g., excitation-inhibition balance^12^) and thus can be validated in experiments^13–15^. However, hand-crafted circuit models come short of capturing the complexity and heterogeneity of single-neuron responses in the cortex.

Single neurons in areas such as the prefrontal cortex (PFC) show heterogeneous tuning to multiple task variables^16–21^. Similar responses arise in recurrent neural network (RNN) models optimized to perform cognitive tasks^19, 22–24^. Yet, the complexity of the RNN activity and connectivity obscure the interpretation of circuit mechanisms in these networks. Heterogeneous neural responses are usually analyzed with dimensionality reduction methods to reveal how latent behavioral variables become mixed within neural populations^21^, ^25–27^. These methods seek low-dimensional projections of neural activity which correlate with specific task variables (Fig. 1a,b). However, these correlation based methods are not constrained by any explicit mechanistic hypotheses. Therefore, it remains unclear whether the inferred latent variables have behavioral significance and how solutions found by RNNs relate to mechanisms posed by circuit models.

**Figure 1.**
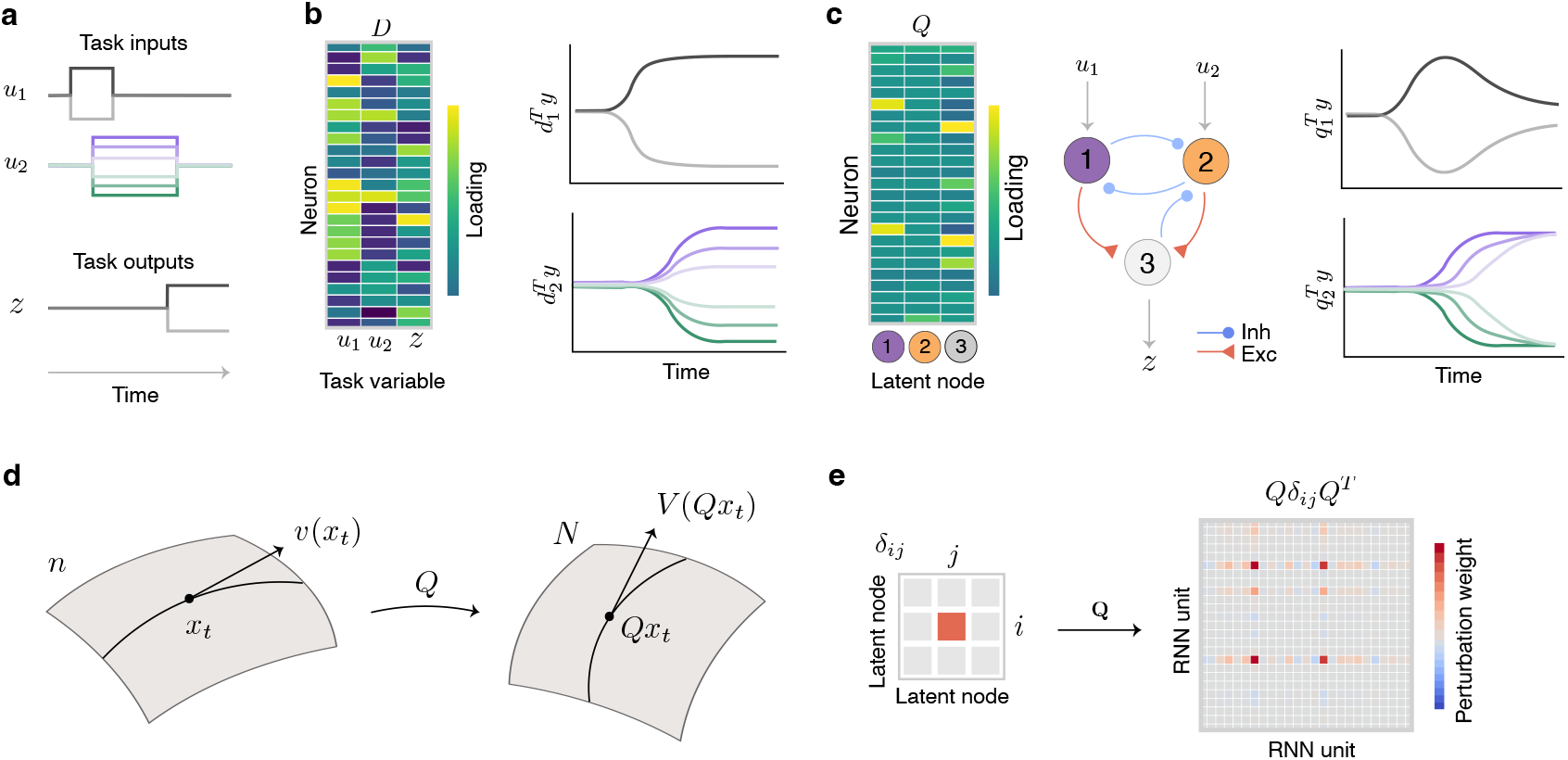
Latent circuit model of heterogeneous neural responses during cognitive tasks. (**a**) A cognitive task requires producing desired behavioral outputs *z* prompted by external inputs *u*. The inputs *u* and outputs *z* are the task variables. (**b**) Dimensionality reduction based on correlation between neural activity and task variables. Decoding matrix *D* defines a projection from neural activity space onto task variables (*left*). Each column of *D* defines an axis in state space such that the projection of neural activity onto this axis correlates with a specific task variable (*right*). (**c**) Latent circuit model. Embedding matrix *Q* defines a projection from neural activity space onto nodes of the latent circuit (*left*). The nodes interact through recurrent dynamics Eq. 2, are driven by task inputs *u*, and generate task outputs *z* (*center*). Each column of *Q* defines an axis in state space such that the projection of neural activity onto this axis correlates with the activity of a node in the latent circuit (*right*). (**d**) Mapping of trajectories gives rise to mapping of vector fields between the latent circuit model and high-dimensional system. (**e**) The relationship between connectivity of the latent circuit and RNN enables us to translate connectivity perturbations. Perturbing connection *δ_ij_* from node *j* to node *i* in the latent circuit maps onto patterned connectivity perturbation 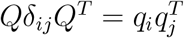 in the RNN.

The glaring gap between these two perspectives is apparent in recent studies of the PFC role in context-dependent decision making. Context-dependent decision making requires flexible trial-by-trial switching between alternative stimulus-response mappings. Most circuit models hypothesize a relatively simple mechanism based on inhibition of sensory representations irrelevant in the current context^8–10, 28, 29^. In contrast, dimensionality reduction methods applied to PFC data or RNN activity show minimal suppression of irrelevant sensory responses^19^, ^21^, seemingly invalidating the inhibitory circuit mechanism. However, since dimensionality reduction methods bear no links to the underlying network connectivity, whether the two perspectives are incompatible remains an open question.

We bridge these two perspectives by developing a modeling framework at the intersection of dimensionality reduction and network connectivity inference. Our framework infers a low-dimensional connectivity structure—the latent circuit—from high-dimensional neural response data. The latent circuit captures mechanistic interactions between task variables and their heterogeneous mixing in single-neuron responses. We applied latent circuit inference to RNNs optimized on a context-dependent decisionmaking task and found a circuit mechanism based on inhibition of irrelevant sensory representations. We validated this mechanism by confirming the behavioral effects of patterned perturbations of the RNN activity and connectivity predicted by the latent circuit model. In contrast, a linear decoder fitted to RNN activity identified a subspace where irrelevant sensory representations were not suppressed, but perturbations within this subspace had no behavioral effects. Our latent circuit framework enables mechanistic interpretation of heterogeneous neural responses and opens the possibility of causal testing of inferred mechanisms in perturbation experiments.

## Results

### Latent circuit model

To bridge the gap between dimensionality reduction, circuit mechanisms and single-neuron heterogeneity, we develop a latent circuit model (Fig. 1c). Similar to other dimensionality reduction methods, we model high-dimensional neural responses *y* ∈ ℝ*^N^* (*N* is the number of neurons) during a cognitive task using low-dimensional latent variables *x* ∈ ℝ*^n^* as:

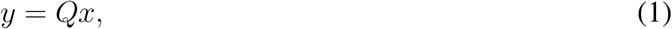

where *Q* ∈ ℝ*^N×n^* is an orthonormal embedding matrix and *n* ≪ *N*. The latent variables *x* in our model are constrained to be nodes in a neural circuit with dynamics

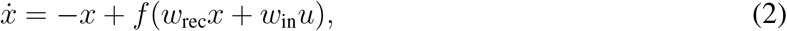

where *f* is a non-linear activation function. The choice of non-linearity *f* presents a tradeoff between model complexity and interpretability. We choose *f* to be a rectified linear function. The latent nodes in the circuit interact via the recurrent connectivity *w*_rec_ and receive external task inputs *u* through the input connectivity *w*_in_. We also require the latent circuit to implement the behavioral task, i.e. we can read out the task outputs *z* from the circuit activity via the output connectivity *w*_out_:

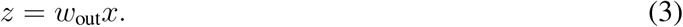

The latent circuit model captures task-related neural activity in the low-dimensional subspace spanned by the columns of *Q*, with dynamics within this subspace generated by the neural circuit Eq. 2.

In the latent circuit model, the heterogeneity of single-neuron responses has three possible sources: mixing of task inputs to the latent circuit via *w*_in_, recurrent interactions between latent nodes via *w*_rec_, and linear mixing of representations in single neurons via the embedding *Q*. The orthonormality constraint on *Q* has a geometric interpretation for the latent circuit embedding, that is we only allow rigid embeddings which rotate the circuit but do not deform it. This constraint implies that the projection defined by the transpose matrix *Q^T^* is a dimensionality reduction, in which projection onto the *i*-th column of *Q* correlates with the activity of the *i*-th node in the latent circuit. Conversely, the image of each latent node *i* is a high-dimensional activity pattern given by the column *q_i_* of the matrix *Q*. Thus, the latent circuit provides a dimensionality reduction which incorporates an explicit mechanistic hypothesis for how the resulting low-dimensional dynamics are generated.

We can infer the latent circuit from the high-dimensional neural activity *y* by minimizing the loss function 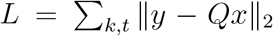, where *k* and *t* index trials and time within a trial, respectively (Methods). The latent circuit model is therefore a linear regression in which the predictor variables arise as the activity of nodes in a non-linear neural circuit. Accordingly, the minimization of *L* is a non-linear least-squares optimization problem^30^, in which we simultaneously search for a projection of the high-dimensional activity and a low-dimensional neural circuit which generates dynamics in the projection. We find the latent circuit parameters *Q*, *w*_rec_, *w*_in_, and *w*_out_ using stochastic gradient descent and backpropagation through time.

In general, it is not obvious under what circumstances we can satisfactorily fit a latent circuit model to the responses of a high-dimensional system. If, for example, solutions to cognitive tasks that emerge in large systems are qualitatively different from mechanisms operating in small circuits, then we should not be able to adequately fit task-related dynamics of the large system with a low-dimensional circuit model. However, the existence of a low-dimensional circuit solution that accurately captures dynamics of the large system would suggest that this circuit mechanism may be latent in the high-dimensional system.

### Interpreting latent connectivity

The advantage of the mechanistic model for latent dynamics is that we can interpret the latent connectivity and relate it to the connectivity of the high-dimensional system. In this context, RNNs optimized to perform a cognitive task provide an ideal setting for testing and validating the latent circuit inference. RNNs mimic the heterogeneity and mixed selectivity of neural responses in the cortex during cognitive tasks, while providing a full access to each unit’s activity, network connectivity, and behavioral outcomes.

To interpret the latent connectivity, we differentiate the embedding Eq. 1 to obtain the correspondence between vector fields of the high-dimensional and low-dimensional dynamical systems (Fig. 1d, Methods):

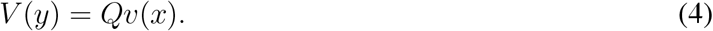

Here 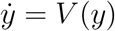 is the vector field of the high-dimensional system, and 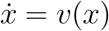 is the vector field of the latent circuit (Eq. 2). This equation states that the subspace spanned by the columns of *Q* is an invariant subspace of the high-dimensional system, i.e. the vector field at any point in this subspace lies entirely in this subspace. Using the orthonormality of *Q*, we then derive the relationship:

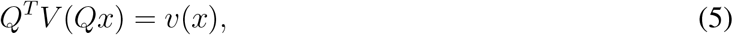

which asserts that the latent vector field *v*(*x*) describes dynamics of the high-dimensional system in this invariant subspace.

For RNNs, we can further derive an explicit relationship between connectivity matrices of the high-dimensional network and low-dimensional latent circuit. We substitute the dynamical equations for the RNN and latent circuit into Eq. 5 and linearize near the origin to obtain (Methods):

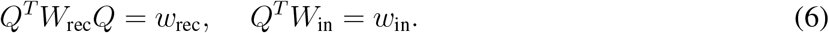

Here *W*_rec_ and *W*_in_ are the recurrent and input connectivity matrices in the RNN, respectively (Methods). This relationship between connectivity matrices relies on the form of dynamical equations for the RNN and latent circuit, in which the connectivity gives a linear approximation of the vector field near the origin. Although we obtained these equalities by linearizing dynamics near the origin, Eq. 6 holds in general, because it is a statement about fixed connectivity structures of a non-linear RNN and the latent circuit.

The relation Eq. 6 shows that the latent circuit connectivity *w*_rec_ can be viewed as a low-dimensional structure in the connectivity of high-dimensional network, which captures interactions between the latent variables defined by the columns of *Q*. The form of Eq. 6 does not necessarily imply that the recurrent connectivity *W*_rec_ is low rank^31^. Rather, it is a weaker condition that the linear subspace defined by *Q* is invariant for the high-dimensional network. In practice, we search for the latent circuit by minimizing the loss function *L*. If *L* is not exactly equal zero, then Eq. 1 and consequently Eq. 6 hold only approximately.

The relation between connectivity matrices has the powerful consequence that we can validate the latent circuit mechanism directly in the RNN connectivity. First, if the latent circuit faithfully describes the mechanism operating in the RNN, by conjugating the RNN connectivity matrix with *Q* (Eq. 6), we expect to find low-dimensional connectivity structure similar to the latent circuit connectivity. Such an agreement is nontrivial, because the latent circuit inference uses only RNN activity without knowledge of the RNN connectivity. Second, Eq. 6 enables us to translate connectivity perturbations in the latent circuit onto the connectivity perturbations in the RNN. Specifically, a change in the connection *δ_ij_* between nodes *i* and *j* in the latent circuit maps onto a patterned perturbation of the RNN connectivity matrix (Fig. 1e, Methods):

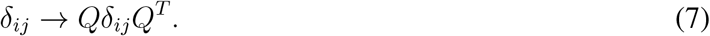

Since the latent circuit is an interpretable mechanistic model of the cognitive task, it inherits the ability of traditional neural circuit models to predict interpretable changes in behavioral performance under specific perturbations of the circuit structure. By translating latent connectivity perturbations onto the RNN, we can verify whether these connectivity perturbations affect RNN behavioral performance as predicted by the latent circuit model. Such verification can ultimately validate the latent circuit mechanism directly in the RNN by bridging all the way from RNN connectivity to behavioral performance. The validation of inferred circuit mechanisms via RNN perturbations is critical, because the fit quality alone does not guarantee that the inferred model captures the correct mechanism that generated data^32^. Moreover, it is generally uncertain whether interpretable circuit mechanisms exist in RNNs optimized to perform cognitive tasks. Thus, confirming predicted behavioral effects of connectivity perturbations serves as a proxy for ground truth when establishing the existence of the inferred circuit mechanism in the RNN.

### Latent circuit mechanism in RNN model of context-dependent decision making

We applied our latent circuit inference to RNNs optimized to perform a context-dependent decision-making task. Contextdependent behavior requires flexible trial-by-trial switching between alternative stimulus-response mappings. Context-dependent decision-making tasks have been extensively used to study the role of PFC and other cortical areas in cognitive control^29^. We used a specific version of the task, which requires to discriminate either the color or motion feature of a sensory stimulus depending on the context cue^19^ (Fig. 2a). At the beginning of each trial, the context cue briefly appears to indicate either the color or motion context for the current trial. After a short delay, a sensory stimulus appears which consists of motion and color features. The right motion and red color are associated with the right choice, and the left motion and green color with the left choice. The strength of motion and color stimuli varies from trial to trial as quantified by the motion and color coherence. In the color context, the choice should be made according to the color and ignoring motion stimulus, and vice versa in the motion context.

**Figure 2.**
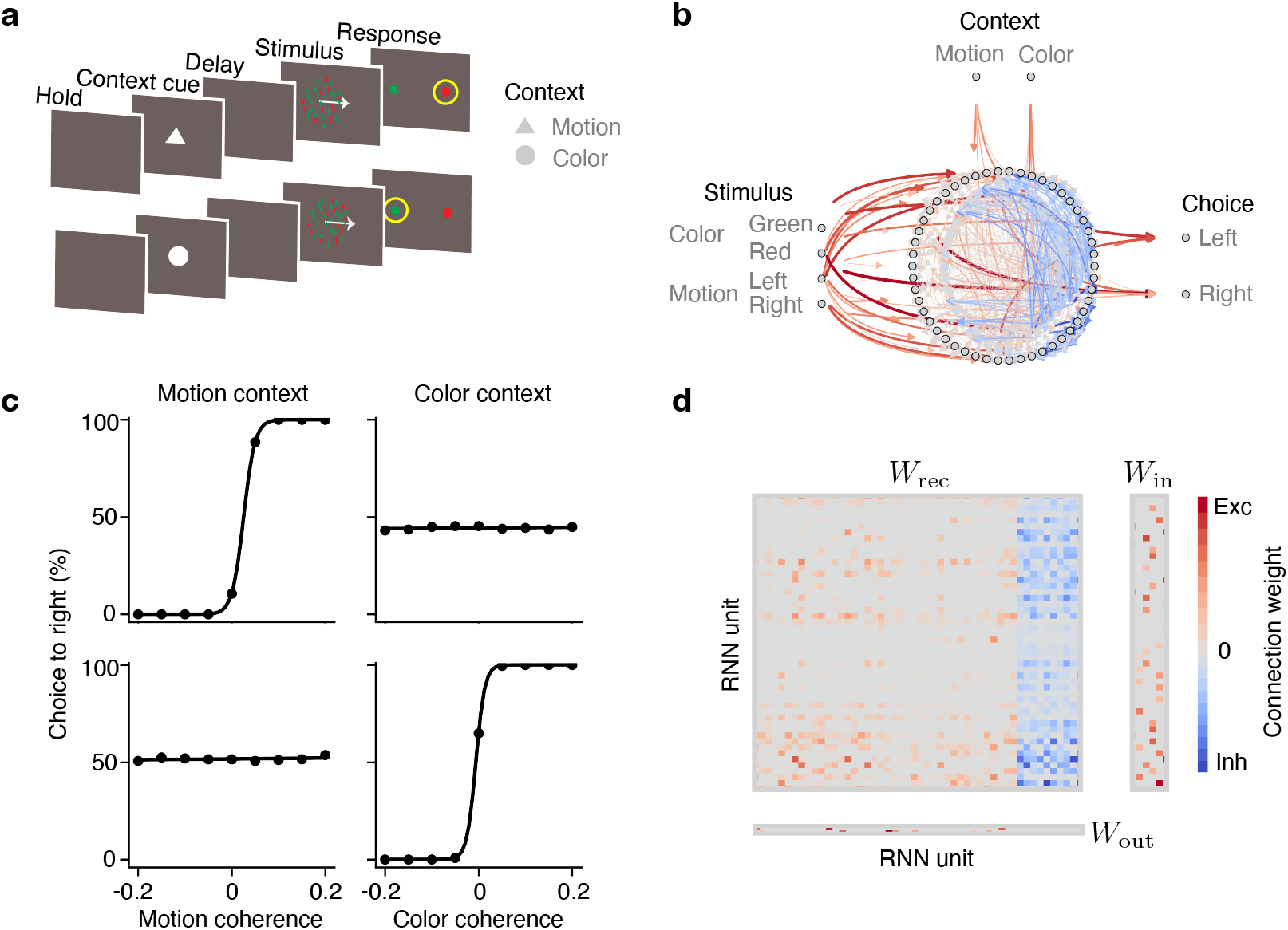
RNN model of context-dependent decision making. (**a**) Context-dependent decision-making task. Each trial begins with a brief baseline period (Hold). Then, a context cue is presented (Context cue). After a short delay (Delay), the color and motion stimuli are presented (Stimulus) and response can be made at any time. The same stimulus can map on different responses depending on the context (Response, yellow circle). (**b**) Architecture of RNN model. Six external inputs *u* drive a recurrent network of 50 units. Choices are readout from the RNN activity at two outputs *z*. RNN is constrained by Dale’s law with 40 excitatory and 10 inhibitory units. (**c**) Psychometric functions show that RNN successfully learns the task: it responds to relevant stimuli and ignores irrelevant stimuli in each context. (**d**) RNN connectivity after training appears complex.

We trained an RNN to perform this context-dependent decision-making task (Fig. 2b, Methods). The RNN consisted of 50 recurrently connected units, 40 excitatory and 10 inhibitory^22^. The RNN received six time-varying inputs *u*: two inputs indicating the color and motion context, and four inputs representing motion (left and right) and color (red and green) stimuli. We trained the RNN to report its decision by elevating one of two outputs *z*, corresponding to the left versus right choice. After training, the psychometric functions confirm that the RNN successfully learns the task, i.e. makes choices according to the relevant stimulus and ignores the irrelevant stimulus in each context (Fig. 2c). Single units in the RNN show heterogeneous mixed selectivity for multiple task variables. The connectivity matrix of the trained RNN appears complex, and the connectivity structure responsible for generating the correct behavioral outputs is not immediately obvious (Fig. 2d).

To reveal the circuit mechanism for context-dependent decision-making in the RNN, we fitted the latent circuit model to the responses of RNN units during the task. The latent circuit model consisted of eight nodes corresponding to the eight task variables: two context nodes, four sensory nodes, and two choice nodes. Each node inherits its identity from the input it receives or the output it sends out, which makes the latent circuit model interpretable. This choice of the latent circuit dimensionality is consistent with the observation that the dimensionality of RNN responses after training is usually close to the total number of inputs and outputs (first eight principal components accounted for 97.9% of total variance in RNN responses). The fitted latent circuit model captured an overwhelming amount of variance in the RNN activity (coefficient of determination *r*^2^ = 0.96 on test data).

The inferred recurrent connectivity *w*_rec_ of the latent circuit revealed an interpretable mechanism for context-dependent decision making (Fig. 3a,b). In the latent circuit, sensory nodes representing stimuli associated with the left choice (left motion and green color) have excitatory connections to the left choice node, and sensory nodes representing stimuli associated with the right choice (right motion and red color) have excitatory connections to the right choice node. This pattern of connections from sensory to choice nodes implements the alternative stimulus-response mappings in the task. Further, the color context node has inhibitory connections to the sensory nodes representing motion, and the motion context node has inhibitory connections to sensory nodes representing color. This pattern of connections from the context to sensory nodes implements a suppression mechanism which inhibits the irrelevant stimulus-response mapping in each context. Since the irrelevant sensory representation is suppressed, it does not drive the decision output. This suppression mechanism based on inhibition of irrelevant representations is qualitatively similar to mechanisms for context-dependent decision-making hypothesized in previous hand-crafted neural circuit models^8^, ^9^.

**Figure 3.**
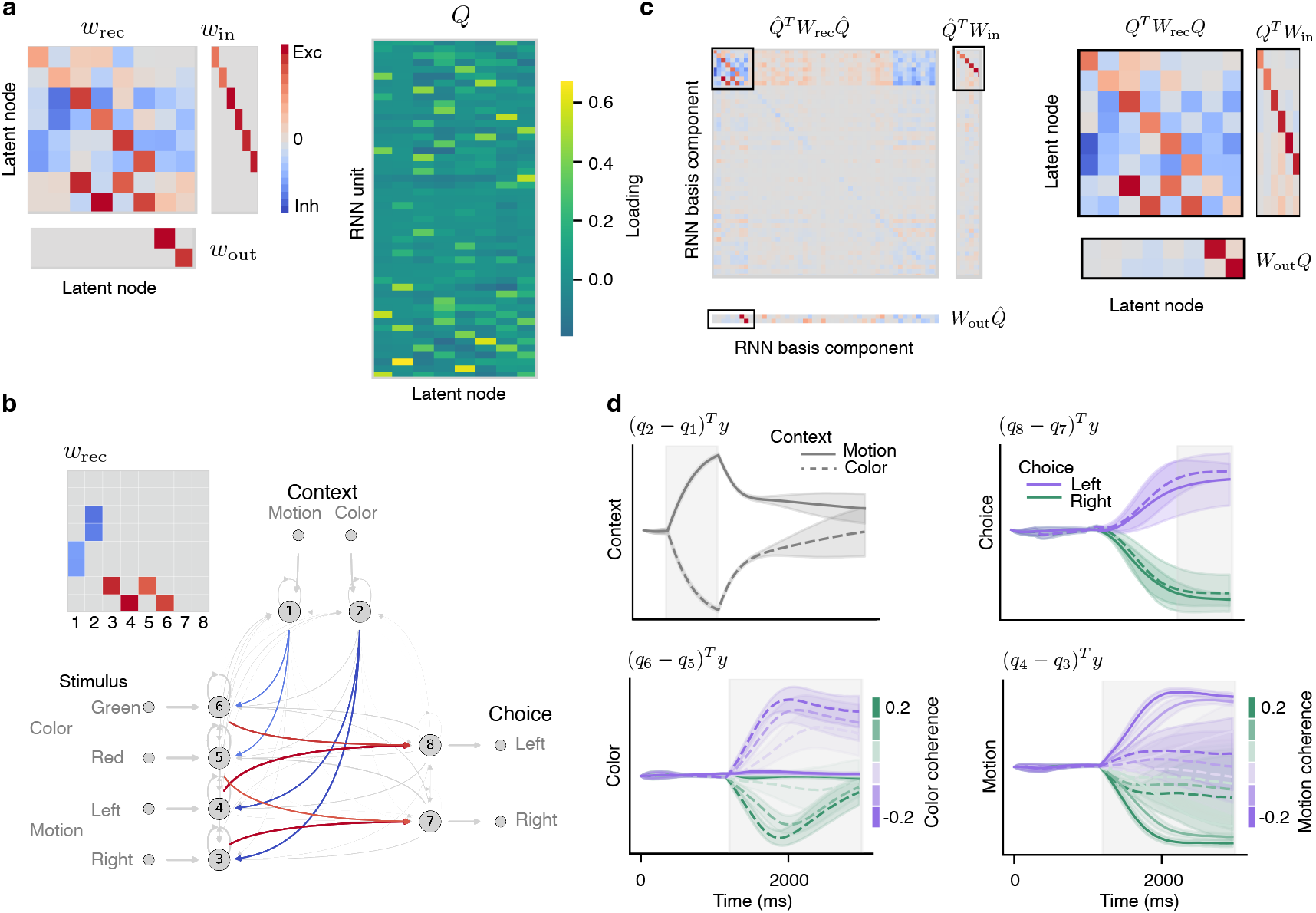
Latent circuit mechanism in RNN performing a context-dependent decision-making task. (**a**) Connectivity matrices of the latent circuit and the embedding matrix *Q* inferred from the responses of RNN performing the context-dependent decision-making task. (**b**) The recurrent connectivity *w*_rec_ in the latent circuit reveals an inhibitory mechanism for context-dependent decision making. The pattern of excitatory connections from sensory to choice nodes implements the alternative stimulus-response mappings (red arrows in the circuit diagram, red squares in the connectivity matrix). The pattern of inhibitory connections from the context to sensory nodes implements a suppression mechanism which inhibits the irrelevant stimulus-response mapping in each context (blue arrows in the circuit diagram, blue squares in the connectivity matrix). The schematic of the connectivity matrix (*upper left*) shows only the eight key connections for clarity. The circuit diagram depicts the full latent circuit connectivity in a. (**c**) We extend *Q* to an orthonormal basis 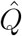 for ℝ*^N^* and transform the RNN connectivity into this basis 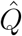 (*left*). The submatrices corresponding to the first *n* = 8 rows and columns (black rectangles, enlarged in *right* panel) closely match the latent circuit connectivity in a (correlation coefficient *r* = 0.89). (**d**) Projections of RNN responses onto low-dimensional subspace defined by the columns of embedding *Q*. By construction, the activity along each projection correlates with the activity difference of two nodes in the latent circuit. Projections onto axes corresponding to motion and color nodes reveal suppression of irrelevant stimulus representations. The grey shading indicates the duration of cue (context axis) and sensory stimulus (motion and color axis) presentation, and response period (choice axis).

We verified whether signatures of the suppression mechanism are evident in the RNN activity. We projected RNN responses onto the columns of *Q*, which are representations of the latent nodes in the RNN (Fig. 3d). These projections reveal RNN activity patterns that are correlated with the activity of nodes in the latent circuit. First, by projecting RNN responses onto the difference of two columns of *Q* corresponding to the context nodes, we obtain a one-dimensional latent variable correlated with the activity difference of the motion-context and color-context nodes in the latent circuit. This projection shows RNN trajectories diverging into opposite directions in state space according to context immediately after onset of the context cue. The activity along this axis then slowly decays, persisting through the stimulus duration. Next, the choice axis is the difference of two columns of *Q* corresponding to the left and right choice nodes in the latent circuit. Projection of the RNN activity onto the choice axis shows trajectories separating according to choice regardless of context. Finally, the motion axis is the difference of columns of *Q* corresponding to the left and right motion nodes, and the color axis is the difference of columns of *Q* corresponding to the red and green color nodes. Projections of the RNN activity onto the motion and color axes reveal representations of relevant sensory stimuli, while representations of irrelevant stimuli are suppressed. In particular, along the color axis, RNN trajectories separate according to color coherence only on color context trials, whereas on motion context trials, the activity along this axis is suppressed. Similarly, activity along the motion axis is suppressed on color context trails. The persistence of context representations and suppression of irrelevant sensory representations in the RNN activity are consistent with the inhibitory mechanism revealed in the latent circuit connectivity *w*_rec_.

We then used the connectivity relationships Eqs. 6,7 to directly validate the inferred latent circuit mechanism in the RNN. We conjugate the RNN connectivity matrix with the embedding matrix *Q*. The resulting connectivity matrices closely match the connectivity structure in the latent circuit (Fig. 3c, correlation coefficient *r* = 0.89). This agreement confirms that the latent connectivity structure indeed exists in the RNN.

To further validate that this latent connectivity structure supports the behavioral task performance, we tested whether patterned perturbations of the RNN connectivity (Eq. 7) produced the same behavioral effects as predicted by the latent circuit model. We consider two perturbations designed to test the inhibitory mechanism suggested by the latent circuit model. The first perturbation corresponds to “turning off” the context mechanism by weakening the inhibitory connections from a context node to sensory nodes representing irrelevant stimuli in that context (Fig. 4a). In the RNN, this perturbation maps onto a patterned change in the recurrent connectivity (Fig. 4b). From interpretability of the latent circuit model, we expect that weakening the inhibitory connections from the motion-context node to sensory nodes representing color should make the circuit sensitive to irrelevant color information on motion context trials. Indeed, weakening these connections in the latent circuit produced the predicted behavioral effect in the psychometric function, visible as a rotation of the decision boundary on motion context trials (Fig. 4a). Perturbations of the RNN connectivity along the corresponding pattern produced similar behavioral effects (Fig. 4b), thus confirming that this connectivity pattern implements suppression of irrelevant sensory representations in the RNN.

**Figure 4.**
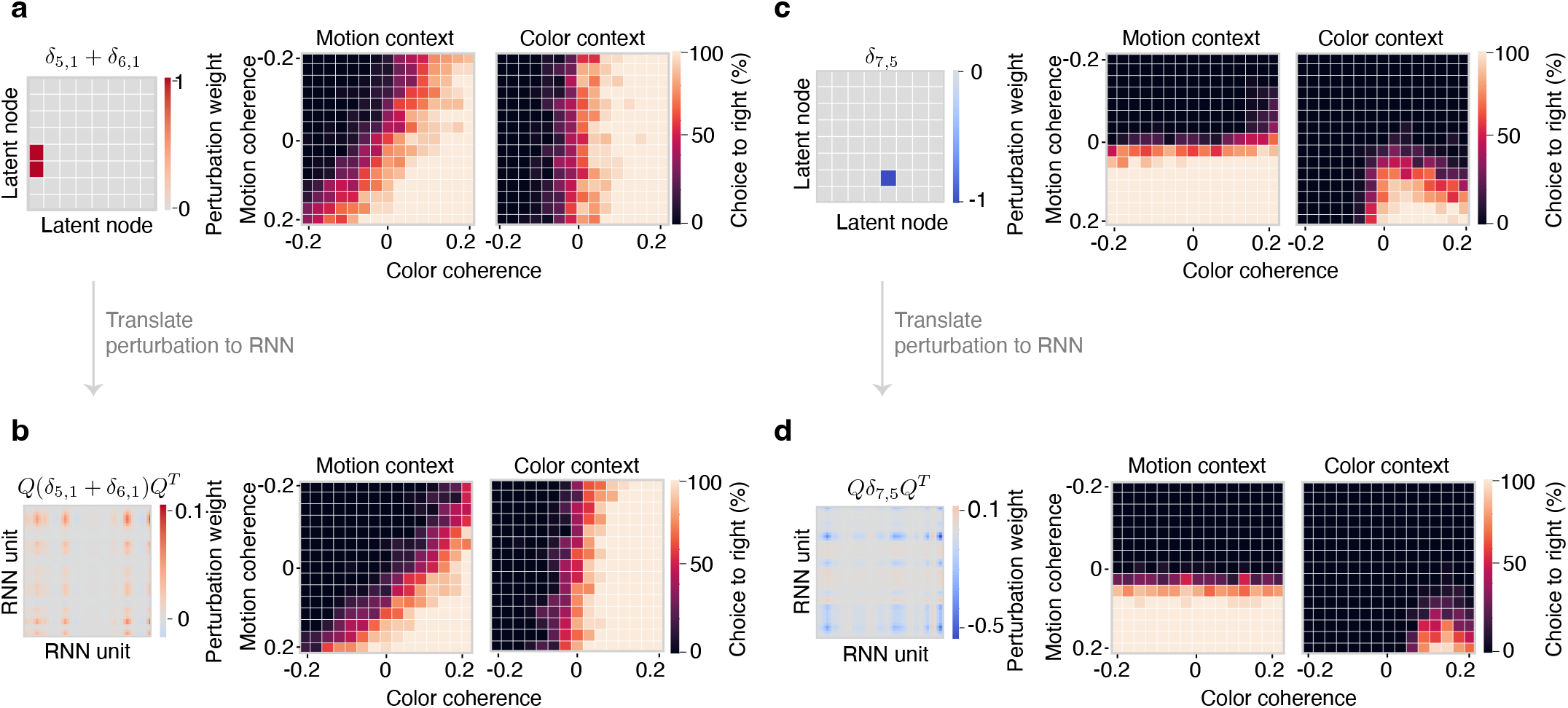
Validating circuit mechanism via perturbations of RNN connectivity. (**a**) Perturbation of the latent circuit connectivity that weakens the inhibitory connection from the motion context node to the sensory nodes representing color (*left*). This perturbation affects behavior making the latent circuit sensitive to irrelevant color information, which is visible as a rotation of the decision boundary on motion context trials in the psychometric function (*right*). (**b**) The perturbation in a of the latent circuit connectivity maps onto patterned connectivity perturbation in RNN (*left*). This perturbation affects the RNN psychometric function as predicted by the latent circuit model (*right*). (**c**) Perturbation of the latent circuit connectivity that weakens the excitatory connection from the node representing red color to the right-choice node (*left*). The effect of this perturbation on behavior is a decrease in the frequency of right choices on color context trials. (**d**) Translation of the latent circuit perturbation in c onto patterned perturbation of the RNN connectivity (*left*) confirms the predicted behavioral effect in the RNN.

The second perturbation corresponds to “turning off” one of the stimulus-response mappings by weakening the excitatory connection from a sensory node to the choice node. From the interpretable latent circuit model, we expect that weakening the excitatory connection from the red color node to the right choice node (Fig. 4c) should impair the network’s ability to make right choices on the color context trials. Weakening this connection in the latent circuit indeed decreased the frequency of right choices on color context trials (Fig. 4c). This perturbation maps onto a patterned connectivity perturbation in the RNN, which produced similar behavioral effects as predicted by the latent circuit (Fig. 4d). This result confirms both the behavioral relevance of the latent sensory representation and the excitatory mechanism by which it drives choices in the RNN.

Together, these results confirm that the RNN utilizes the suppression mechanism in which persistent contextual representations inhibit irrelevant sensory representations. This mechanism is reflected in the low-dimensional dynamics revealed by projecting RNN activity onto axes defined by the embedding *Q*. We identified this mechanism as latent low-dimensional structure in the RNN connectivity and ultimately validated it by confirming behavioral effects of the RNN connectivity perturbations.

### Representations of irrelevant stimuli

Our finding that the RNN uses the inhibitory mechanism for context-dependent decision-making appears in conflict with previous work, which suggested that in both PFC and RNNs, irrelevant sensory responses are not significantly suppressed^19, 21, 33^. This conclusion was derived using dimensionality reduction methods which extract low-dimensional projections that best correlate with task variables (Fig. 1a). However, because these dimensionality reduction methods do not incorporate any explicit mechanistic constraints, it is unclear whether the absence of suppression in these projections is incompatible with the inhibitory mechanism we identified.

To answer this question, we compared projections of the RNN activity onto axes obtained from the latent circuit model and a linear decoder. We trained a linear decoder to predict the signed motion coherence on each trial from the RNN activity (Methods). The decoding weights provide an axis in the RNN state space such that a projection onto this axis correlates with the motion coherence. By projecting RNN responses onto the decoder axis, we find a strong representation of irrelevant motion stimulus on color context trials without noticeable suppression (Fig. 5a). Thus, in our RNN, irrelevant sensory representations appear not suppressed along the decoder axis, whereas they appear suppressed along the axis obtained from the latent circuit model (Fig. 5b). This difference arises from the distinct objectives of each projection: an axis in which stimulus activity is suppressed on half the trials does not meet the objective of the decoder which tries to maximize the stimulus decoding accuracy across all trials. Thus, the linear decoder is biased toward an axis which does not show suppression, providing a misleading picture of dynamics. In principle, we could account for this bias by fitting a linear decoder to restricted data from trials on which the stimulus is relevant, but this procedure would presume knowledge of the mechanism which we are trying to discover in the first place.

**Figure 5.**
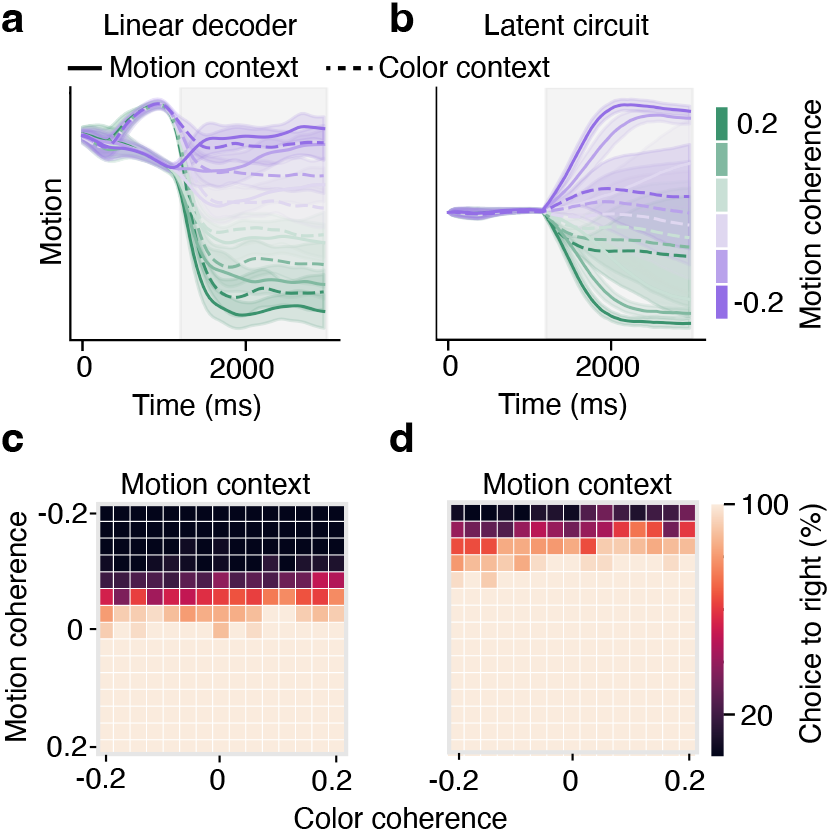
Representation of irrelevant stimuli in RNN does not drive behavior. (**a**) Projection of the RNN responses onto decoder axis reveals representation of motion coherence on both color context and motion context trials. (**b**) Projection of the RNN responses onto motion axis from the latent circuit model reveals that representation of motion coherence is suppressed on color context trials. (**c**) Patterned stimulation of the RNN activity along the decoder axis has little effect on the psychometric function. (**d**) Patterned stimulation of the RNN activity along the motion axis from the latent circuit shifts the decision boundary in the psychometric function consistent with enhanced representation of right motion stimulus. These perturbations reveal that the decoder compromises behavioral relevance for decoding accuracy.

How can we reconcile these qualitatively distinct perspectives on representations of irrelevant stimuli within a single RNN? We hypothesized that the appearance of irrelevant stimulus representations is possible because the linear decoder compromises behavioral relevance and variance explained for decoding accuracy. To test this idea, we stimulated RNN units with activity patterns aligned with the axes obtained from the linear decoder and the latent circuit model. If the corresponding activity patterns are behaviorally relevant, we expect the stimulation to have a significant effect on psychometric functions. Specifically, stimulating the representation of the right motion stimulus should increase the proportion of right choices, shifting the decision boundary on motion context trials. As expected, driving RNN activity along the motion axis of the latent circuit model shifted the decision boundary on motion context trials (Fig. 5d). In contrast, stimulation of the same magnitude along the decoder axis had little effect on the psychometric function (Fig. 5c). This result suggests that the decoder rotates its axis away from the behaviorally relevant latent circuit axis to achieve better decoding of motion coherence across all trials. The irrelevant stimulus representations exist along the decoder axis but do not drive the behavioral output. For context-dependent decision-making, we conclude that the dynamics revealed by the decoder have little behavioral significance and thus do not invalidate the inhibitory mechanism identified by the latent circuit model.

The projection axes obtained via our latent circuit model do not necessarily “demix” task variables. These projections inherit their interpretation from the corresponding nodes in the latent circuit, and nodes are defined by both by their external inputs and recurrent interactions with other nodes. For example, both motion and color axes are modulated by sensory input as well as input from the context nodes. In contrast, other dimensionality reductions methods seek low-dimensional projections that decode individual task variables in orthogonal dimensions, i.e. demix task variables^19, 21, 26^. Our results indicate that “demixing” may be not the right objective for identifying behaviorally relevant patterns in the heterogeneous neural activity.

### Space of latent circuit mechanisms

We next asked whether RNNs optimized to perform the same task can arrive at different latent circuit mechanisms for context-dependent decision-making. We trained an ensemble of 550 RNNs with randomly initialized connectivity to the same level of task performance. After training, there was little variation in behavioral accuracy across these networks (*r*^2^ = 0.92 ± 0.01, coefficient of determination for the RNN and target outputs, mean±s.d.). For each of these RNNs, we fitted an ensemble of latent circuit models starting with random initializations of the latent connectivity parameters and embedding *Q*. To analyze the space of inferred latent circuit mechanisms, we examined the variability within the set of latent recurrent connectivity matrices *w*_rec_. We applied the principal component analysis (PCA) to the flattened connectivity matrices and projected the data onto the first two principal components (Fig. 6a). We found that the set of latent circuit mechanisms lies on a curved one-dimensional manifold with three major clusters. Specifically, the majority of circuits fall within a single cluster, while relatively fewer circuits fall within one of two other clusters branching off of the main cluster. Moreover, the ensemble of latent circuit solutions fitted to responses of a single RNN fall within a close proximity of each other, which indicates the uniqueness of the latent circuit mechanism in each particular network.

**Figure 6.**
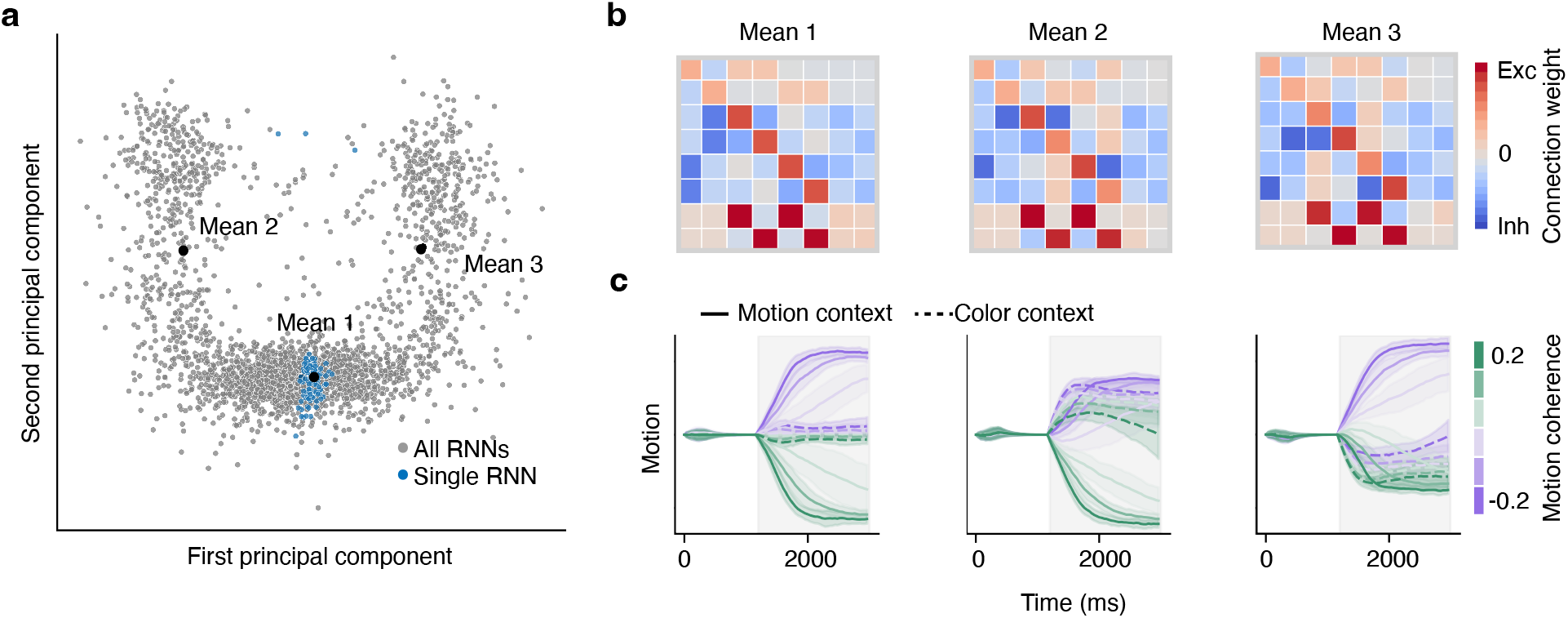
Space of latent circuit mechanisms for context-dependent decision-making in RNNs. (**a**) Projection onto the first two principal components of the set of recurrent connectivity matrices in latent circuits obtained from 550 RNNs performing the context-dependent decision-making task. For each of these RNNs, we fitted an ensemble of latent circuits with random initializations. Latent circuits fitted to responses of a single RNN fall within a close proximity of each other (blue dots). The projection reveals an approximately one-dimensional manifold of latent circuit solutions with three major clusters. The means of these clusters are estimated by fitting a Gaussian mixture model with three components. (**b**) Recurrent connectivity matrices corresponding to the means of three clusters in a. (**c**) Dynamics of the activity difference between two nodes representing motion stimulus in latent circuits with the mean connectivity matrices in b. These dynamics reveal asymmetric representations of stimuli at the extremes of the manifold of latent circuit solutions.

To understand how the circuits vary within this manifold, we computed the mean latent connectivity matrix for each cluster by fitting data with a Gaussian mixture model with three components. The resulting mean connectivity matrices reveal a continuum of circuits that all show signatures of the suppression mechanism in which context nodes inhibit irrelevant sensory nodes (Fig. 6b). In the main cluster, the circuits are balanced and symmetric, with approximately equal strength of excitation or inhibition between nodes representing different contexts and stimulus-response mappings. This balance is reflected in dynamics and the representations of stimuli (Fig. 6c). In the two other clusters, the circuits show asymmetry in the connectivity, with stronger inhibition from context to some sensory nodes, counterbalanced by stronger self-excitation for these sensory nodes. These assymetries are consistently reflected in dynamics and the representations of stimuli, which show a bias toward the left or right stimulus representations depending on the cluster. Although these extreme solutions exploit assymetries in the representations of sensory evidence, they still operate by an inhibitory mechanism in which irrelevant responses are suppressed. Thus, the space of latent circuit solutions found by RNN models of the context-dependent decision-making task can be characterized by a common suppression mechanism.

## Discussion

Single neurons in higher cortical areas, in particular prefrontal cortex, show heterogeneous responses during cognitive tasks, with complex mixed selectivity for multiple sensory, contextual, and motor variables. We developed a latent circuit model which accounts for single-neuron heterogeneity via dimensionality reduction that incorporates explicit neural circuit dynamics in its latent variables. The latent circuit can be interpreted as a low-dimensional connectivity structure that supports the behavioral task performance in a high-dimensional system. This latent connectivity structure can be inferred from neural responses and causally tested via connectivity perturbations in the high-dimensional system.

Previous work highlighted the importance of incorporating dynamics into dimensionality reduction methods^34–38^. However, these methods do not provide a supervised approach for understanding encoding and transformation of task variables and are only rarely constrained by behavioral performance^33^, ^39^. In contrast, the latent circuit model infers an explicit mechanistic hypothesis for how task variables interact to drive behavioral outputs. Additionally, there is extensive work on inferring low-dimensional effective circuits from response data using dynamic causual models (DCMs)^40–42^. Unlike DCMs, our latent circuit model simultaneously infers the dynamics and embedding of the low-dimensional circuit, thus bridging the gap between dimensionality reduction, circuit mechanisms, and single-cell heterogeneity.

Our results have important implications for inference approaches which aim to reconstruct connectivity from neural response data^43, 44^. These methods require large volumes of data to recover full connectivity in large networks. We show that if the responses arise from low-dimensional behavior, then what can be recovered is not the full connectivity, but a latent low-dimensional connectivity structure capturing how the network acts on a low-dimensional activity subspace. From this perspective, a latent circuit model is a framework for inferring low-dimensional latent connectivity. This perspective implies that our approach does not provide a way of constructing high-dimensional models with low-rank solutions^31^, because it is not possible to uniquely recover the high-dimensional connectivity from the low-dimensional latent connectivity structure which is inferred from low-dimensional neural responses.

Using a latent circuit model, we found an interpretable circuit mechanism in RNNs performing a context-dependent decision-making task. The inferred latent circuit connectivity revealed a suppression mechanism in which context nodes inhibit irrelevant sensory nodes. This latent connectivity structure is consistent with low-dimensional dynamics observed in the projections of RNN activity onto the low-dimensional subspace spanned by the embedding *Q*. In these projections, stimulus representations are suppressed when they are irrelevant. The inhibitory mechanism exist as a latent low-dimensional connectivity structure in the RNN, and we causally validated this mechanism via perturbations of the RNN connectivity. The inhibitory mechanism revealed by the latent circuit model is qualitatively similar to previous neural circuit models of how prefrontal cortex flexibly switches between alternative stimulus-response mappings^8–10, 28, 29^. We find that RNNs do not find qualitatively distinct solutions to this task and that complex selectivity of single neurons has a simple explanation as a linear mixing of the low-dimensional latent inhibitory circuit mechanism. Our results suggest that similar interpretable mechanisms may underlie the complex responses of prefrontal neurons during context-dependent behavior.

In contrast to the latent circuit model, dimensionality reduction methods which are not constrained by an explicit mechanistic hypothesis found non-suppressed representations of irrelevant stimuli in PFC and RNNs during context-dependent decision-making^19, 21^. We show that such representations can arise in RNNs which provably implement an inhibitory suppression mechanism. These representations are possible because they exist along dimensions which do not causally drive choices and thus do not affect behavior. Moreover, approaches aiming to decode stimulus information whether or not it is relevant appear to be biased toward finding these behaviorally irrelevant dimensions. Thus, inhibitory mechanisms for cognitive flexibility are in principle compatible with the existence of irrelevant stimulus representations in the prefrontal cortex, and decoding approaches may be biased toward not discovering these mechanisms. Therefore, low-dimensional dynamics obtained by decoding approaches cannot be interpreted as causal mechanisms supporting task behavior.

We find that RNNs do not necessarily find qualitatively new solutions to cognitive tasks than mechanisms hypothesized in hand-crafted neural circuit models. These mechanisms can be found in large RNNs if connectivity is viewed in the appropriate basis. In other words, just as dynamics should be understood in terms of latent variables, it appears connectivity can be understood in terms of interactions between these latent variables. These results agree with recent work on low-rank RNNs^45^ and add to the growing body of literature suggesting that RNNs trained on cognitive tasks do not find qualitatively distinct mechanisms from those which can be constructed by hand^24^, ^46^. We show that these mechanisms can be inferred from the high-dimensional responses using a dimensionality reduction which is constrained by an explicit choice of latent circuit dynamics. Our approach opens new possibilities for causally testing circuit mechanisms supporting complex behavior in perturbation experiments.

## Methods

### Fitting a latent circuit model

We fit the latent circuit model Eqs. 1–3 to neural response data *y* by minimizing the mean squared error loss function:

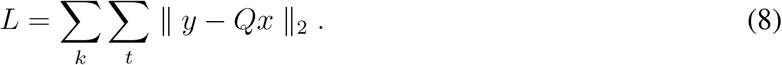

Here *k* indexes the trials and *t* indexes the time within a trial. The orthonormal matrix *Q* is parameterized by

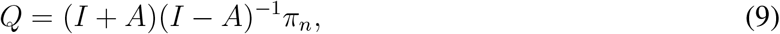

where *π_n_* represents projection onto the first *n* columns. *A* is a skew-symmetric matrix parameterized by

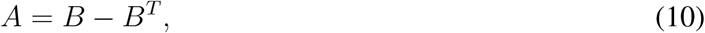

where *B* is an *N* × *n* matrix. At each step of the optimization, we generate a set of trajectories *x* from the latent circuit dynamics and embed these trajectories using the matrix *Q*. The parameters *B*, *w*_rec_, *w*_in_ and *w*_out_ are then updated to minimize *L*. We perform this minimization using PyTorch and the Adam optimizer with default values for the first and second moment estimates, a learning rate of 0.02 and a weight decay of 0.001. We use a minibatch size of 128 trials. We stop the optimization when the loss has not improved by a threshold of 0.001 after a patience of 25 epochs.

We initialize the recurrent matrix *w*_rec_ with zeros, and *w*_in_ with zeros except for positive entries on connections from inputs *u* to their corresponding nodes, and *w*_out_ with zeros except for positive entries on connections from choice nodes to their corresponding outputs *z*. We initialize the entries of matrix *B* from a uniform distribution on [0, 1]. To test for uniqueness of the latent circuit solution in a single RNN, we fit an ensemble of latent circuits with random initializations to the same response data *y*. In this case, we initialize entries in *w*_rec_ from a uniform distribution centered on 0 with standard deviation 1*/n*.

### Relationship between connectivity of RNN and latent circuit

We consider RNNs of the form:

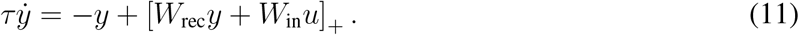

Here [·]_+_ is a rectified linear activation function, *τ* is a time constant, *u* are external task inputs. *W*_rec_ and *W*_in_ are the recurrent and input connectivity matrices, respectively. We read out a set of task outputs *z* from the network activity via the output connectivity matrix *W*_out_:

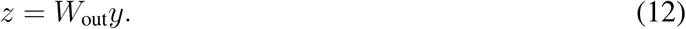

We derive a relationship between the connectivity matrices of the RNN and latent circuit, which allows us to interpret the latent circuit connectivity as a latent connectivity structure in the RNN. To derive this relationship, we differentiate the embedding Eq. 1 with respect to time and obtain the relationship between the vector fields of the RNN and latent circuit:

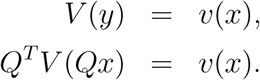

Here the vector fields *V* (*y*) of the RNN and *v*(*x*) of the latent circuit are given by Eq. 11 and Eq. 2, respectively. We then linearize the vector fields in a neighborhood of *x* = 0 and *u* = 0 and obtain:

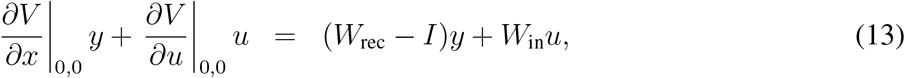

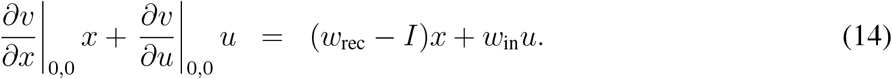

It follows that we have the equality:

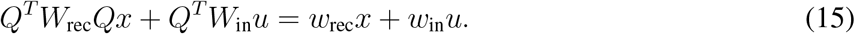

If we assume that this equality holds in some open set, then we can equate terms to obtain:

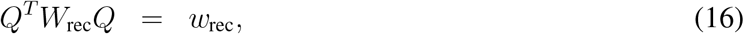

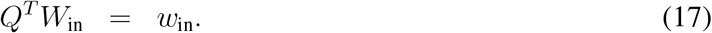

This assumption is likely not fully satisfied in the setting of cognitive tasks, because the sets of inputs *u* and latent states *x* are typically low-dimensional. Therefore, the above equalities may hold only approximately.

To understand how perturbations of connectivity in the latent circuit map onto the RNN, we view perturbations as vectors in the space of matrices. We denote *A ⋅ B* the dot product between the matrices *A* and *B* represented as vectors in the space of matrices, i.e. 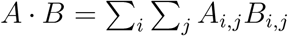. Using Eqs. 16,17, we then translate connectivity perturbations from the latent circuit to the RNN:

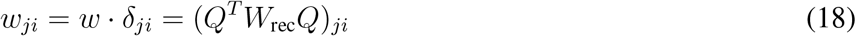

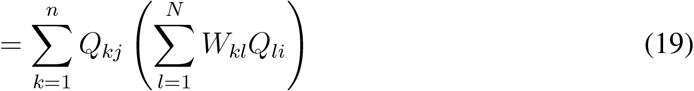

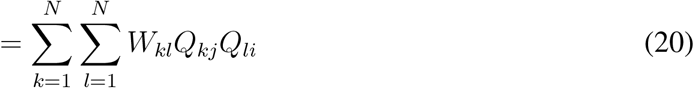

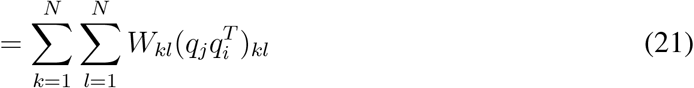

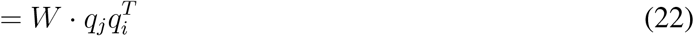

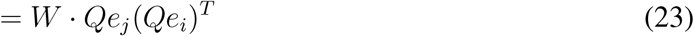

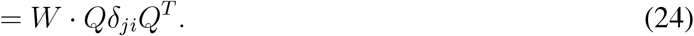

This chain of equalities shows how to translate perturbations of the latent circuit connectivity in the direction *δ_ji_* onto patterned perturbations in the RNN:

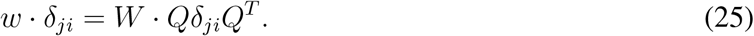

Thus, to perturb the latent connection *w_ji_*, we perturb the matrix *W* in the direction *Qδ_ji_Q^T^*. In other words, to increase the dot product between *W* and *Qδ_ji_Q^T^* in the space of matrices, we add multiples of *Qδ_ji_Q^T^* to *W*. Any perturbation orthogonal to *Qδ_ji_Q^T^* does not change the dot product and hence has no effect on the latent connection *w_ji_*.

### RNN simulations

We simulate dynamics of time-discretized RNNs using the general framework for modeling cognitive tasks^22^. We consider RNNs with positive activity and *N* = 50 recurrent units. We obtained the same results with networks consisting of *N* = 150 units. We discretize the RNN dynamics Eq. 11 using the first-order Euler scheme with a time-step Δ*t* and add a noise term to obtain:

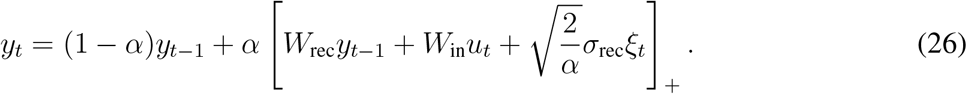

Here *α* = Δ*t/τ* and 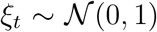 is a random variable sampled from the standard normal distribution. We set the time constant *τ* = 200 ms, the discretization time-step Δ*t* = 40 ms, and the noise magnitude *σ*_rec_ = 0.15. The input and output matrices are constrained to have positive entries. The input matrix is constrained to be orthogonal. The recurrent matrix is constrained to satisfy the Dale’s law with 80% excitatory units and 20% inhibitory units. The RNN simulation and training were implemented in PyTorch and Skorch.

### Context-dependent decision-making task

To model the context-dependent decision-making task, the network receives six inputs *u* corresponding to two context cues (*u*_m_ - motion context, *u*_c_ - color context), and sensory evidence streams for motion (*u*_m,L_ - motion left, *u*_m,R_ - motion right) and color (*u*_c,R_ - color red, *u*_c,G_ - color green). Each trial begins with a presentation of a context cue from *t* = 320 to *t* = 1, 000 ms. On motion context trials, the cue input is set to *u*_m_ = 1.2 and *u*_c_ = 0.2, and vice versa on color context trials. During this epoch, we require that the network does not respond on the outputs by setting *z*_target_ = 0.2. After another delay of 200 ms, so that the network must maintain a memory of the context cue, the inputs corresponding to motion and color sensory evidence are presented at *t* = 1, 200 ms for the remaining duration of the trial. After 2, 250 ms, the targets are defined by *z*_target,1_ = 1.2 and *z*_target,2_ = 0.2 for right choices and vice versa for left choices. The strength of sensory evidence for motion and color varies from trial to trial randomly controlled by the stimulus coherence. We use motion coherence *m*_c_ and color coherence *c*_c_ ranging from −0.2 to 0.2 chosen from the set {−0.2, −0.12, −0.04, 0.04, 0.12, 0.2}. For each coherence level, the motion and color inputs are given by:

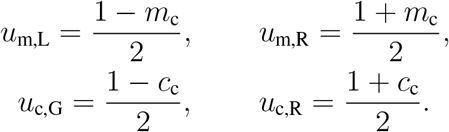

With these definitions, positive motion and color coherence provide evidence for the right choice, and negative motion and color coherence provide evidence for the left choice. At each simulation time step, we add an independent noise term to each of the inputs 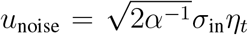, where 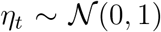 is a random variable sampled from the standard normal distribution. The input noise strength is *σ*_in_ = 0.01. A baseline input *u*_0_ = 0.2 is added to each of the inputs at each time step.

### RNN training

To train the RNN, we minimize the mean squared error between the output *z*(*t*) and the target *z*_target_(*t*):

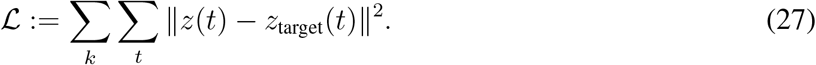

Here *k* is the trial number and *t* is the time step within a trial. To encourage the network to integrate sensory evidence over time, *t* is taken to be greater than 2, 250 ms so that errors are only penalized in the last 750 ms of each trial. The training is performed with the Adam algorithm for stochastic gradient descent. We used the default set of values for the decay rate of the first and second moment estimates, 0.9 and 0.999, respectively. We used a learning rate of 0.01.

The recurrent connection matrix is initialized so that excitatory connections are independent Gaussian random variables with mean 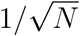 and variance 1*/N*. Inhibitory connections are initialized with mean 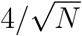 and variance 1*/N*. The matrix is then scaled so that its spectral radius is 1.5. To implement Dale’s law, connections are clipped to zero after each training step if they change sign. During training, we used mini-batches of 128 trials with 1,800 trials total.

To assess performance, a choice for the RNN was defined as the sign of the difference between output units at the end of the trial. Psychometric functions were then computed as the percent of choices to the right for each combination of context, motion coherence and color coherence.

### Linear decoding

To decode motion coherence from RNN responses, we fit a linear regression model:

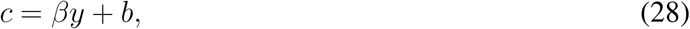

where *β* ∈ ℝ^1*×N*^ is the vector of regression coefficients, *c* ∈ ℝ^1*×k*^ is the motion coherence on each trial, *b* ∈ ℝ is a bias term and *y* ∈ ℝ^*N×k*^ is the RNN responses in the stimulus epoch of each trial. To control for overfitting, we use k-fold cross validation with *k* = 5. We compared the predictive *r*^2^ and their deviation across folds for each of the *k* pairs of training and test data. There was no significant difference between training and test scores, suggesting that the model did not overfit. After fitting, we used the vector of regression coefficients *β* to define the decoder axis on which we project RNN responses.

## Acknowledgements

This work was supported by the Swartz Foundation (C.L.), the NIH grant RF1DA055666 (T.A.E.), and Alfred P. Sloan Foundation Research Fellowship (T.A.E.).

## Author contributions

C.L. and T.A.E. designed the research and developed the framework. C.L. developed the code and performed computer simulations, analysis and calculations. C.L. and T.A.E. wrote the paper.

## Data availability

The synthetic data used in this study can be reproduced using the source code.

## Code availability

The source code to reproduce the results of this study will be made available on GitHub upon publication.

